# Irregular chromatin: packing density, fiber width and occurrence of heterogeneous clusters

**DOI:** 10.1101/453126

**Authors:** Gaurav Bajpai, Ranjith Padinhateeri

## Abstract

How chromatin is folded in the lengthscale of a gene is an open question. Recent experiments have suggested that, *in vivo*, chromatin is folded in an irregular manner and not as an ordered fiber with a width of 30 nm expected from theories of higher order packaging. Using computational methods, we examine how the interplay between DNA-bending non histone proteins, histone tails, intra-chromatin electrostatic and other interactions decide the nature of packaging of chromatin. We show that while the DNA-bending non histone proteins make the chromatin irregular, they may not alter the packing density and size of the fiber. We find that the length of the interacting region and intra-chromatin electrostatic interactions influence the packing density, clustering of nucleosomes, and the width of the chromatin fiber. Our results suggest that the actively maintained heterogeneity in the interaction pattern will play an an important role in deciding the nature of packaging of chromatin.

## Introduction

DNA is a very long polymer that contains the genetic code. Inside biological cells, DNA is not in its bare form; rather it is folded, decorated, and packaged with the help of myriads of proteins and chemical groups into a functional structure known as chromatin (1). The purpose of folding DNA into chromatin is not just for the packaging and storage of the polymer but also for regulating the accessibility and reading of the genetic code. How this one-dimensional(1D) sequence information is folded and packaged into a three-dimensional(3D) chromatin organization is poorly understood. Research in the last few decades has shown that stretches of DNA are wrapped around a multimer complex of histone proteins leading to the formation of an array of nucleosomes (2, 3). However, exactly how this nucleosomal array is further folded and packaged is an open question (4).

In the packaging of chromatin, the nature of interaction and structural details of nucleosomes are thought to play an important role (5, 6). DNA is a negatively charged polymer while histone proteins are predominantly positively charged. In a nucleosome, histone protein subunits (H2A-H2B, H3-H4) have tail-like regions giving rise to a nucleosome complex having multiple tails protruding out of the core region (7). Given that these tails are electrostatically charged, and that various chemical modifications (e.g. acetylation) can alter the charge density, the histone tails and electrostatic interactions are thought to play crucial roles in higher order packaging of chromatin (8–10). Apart from the core histones, the linker histone (H1) is also thought to play an important role in chromatin organization. The H1 protein is known to bind between DNA segments that are entering and exiting a nucleosome and thought to stabilize the nucleosome structure (11–13).

How a string of nucleosomes, known as 10-nm wide chromatin, gets folded further to form a higher order structure is a question that has not been settled even after nearly four decades of research (4, 14). *In vitro* reconstitution of chromatin, based on the array of nucleosomes, suggested that the nucleosomes will form a regular zig zag-like structure having a width of 30 nm (7, 15, 16). Based on other *in vitro* experiments, there is also an alternate hypothesis that the chromatin should be solenoid with 30 nm width (17, 18). Simple polymer physics theories, accounting for certain physical aspects of DNA stiffness and the structure of the nucleosomes, predicted regular 30nm chromatin organization (19–22). The extent of folding of this string of nucleosomes into chromatin is measured by computing the packing density, which is roughly defined as the number of nucleosomes packed in every 11 nm effective length of the chromatin fiber. Analysis of *in vitro* experimental data and simulation results showed that the packing density of 30 nm chromatin fiber can vary from 6 nucleosomes/11 nm to 12 nucleosomes/11 nm depending on different conditions (11, 12, 16, 17, 23–32).

However, most of the recent experiments suggest that, *in vivo*, chromatin does not have any regular structure (33–36). Cryo-electron microscopy (cryo-EM) study on mitotic and interphase cells suggest that there is no 30nm-wide structure *in vivo* (33–35). Recent studies with advanced EM techniques to visualize chromosomes in interphase and mitotic cells suggest that chromatin is folded into an irregular chain having ≈ 5–24 nm wide structures (37). Chromatin conformation capture (Hi-C) experiments suggest that in interphase chromatin is not a homogeneous or a regular structure (38). It is organized into topologically associated domains (TADs) and lamina associated domains (LADs) having open (euchromatin) and compact (heterochromatin) domains distributed across the nucleus and have a fractal nature (38, 39).

There have been many recent theoretical/computational studies trying to understand the irregular nature of chromatin (10, 40– 42). It has been hypothesized that molecular crowding in the cell may lead to the formation of irregular chromatin (34). It has been shown that the variability in linker length and other factors give rise to a polymorphic structure of chromatin (42, 43). Our own earlier work suggested that DNA-bending non histone protein(NHP) can make chromatin irregular (41).

Even though there is a vast literature on chromatin organization, many important questions remain unanswered. Given that chromatin is irregular in the length-scale of genes, the first question is about its compactness. How compact is the irregular chromatin when compared to the regular structures? How does the irregular nature influence the packing density? Given that recent experiments observed chromatin of width 5–24 nm, can we have a theoretical explanation for this range of widths? Given that chromatin is heterogeneous in terms of interaction potentials (e.g., spatial variation of histone modifications), how will this affect chromatin configurations? Does the spatial extent/variation in the modification pattern affect the packing density and width?

In this paper, we address these questions by performing coarse-grained molecular simulations and studying chromatin organization in 3D. We simulate a polymer model of chromatin having nucleosomes with explicit histone tails, electrostatic interactions, and other intra-chromatin interactions. We also account for the spatial variations of the interaction potentials to mimic interaction heterogeneity due to histone modification patterns. We show how DNA-bending NHPs and heterogeneous interactions among nucleosome particles affect packaging of irregular chromatin. Our work suggests why chromatin is less than 30 nm wide even when it is in a packaged heterochromatin state, as observed in recent experiments.

## Model and Method

We present a coarse-grained model where chromatin is modelled as a set of bead-spring chains having five major parts namely (i) the DNA polymer chain, (ii) the core histone, (iii) explicit histone tails, (iv) the linker histone and (v) the DNA-bending non-histone protein (see Fig. 1).

**Figure 1.**
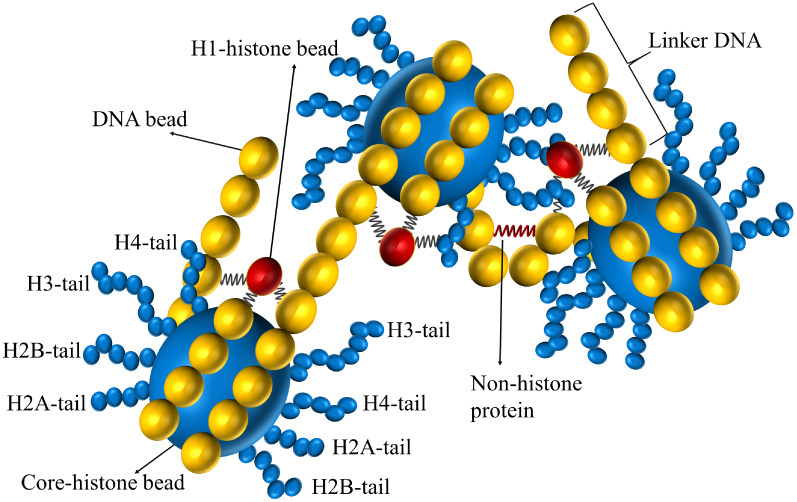
Schematic diagram describing the model: DNA is modeled as a polymer made of type-1 beads(yellow) having a diameter of 3.4 nm. 14 DNA-beads are wrapped around the core histone bead (type-2, big blue bead, 5.25 nm in diameter). Histone tails are modeled as flexible bead-spring polymer chains where each histone-tail bead (type-3, small blue bead) is 1.56 nm in diameter and have the following lengths: tail-1 (H2A)= 4 beads, tail-2 (H2B)= 5 beads, tail-3 (H3)= 8 beads, and tail-4 (H4)=5 beads. H1 histone (red bead, type-4) is connected to three DNA beads— the entry bead, exit bead, and the central bead of the DNA wrapped around the core histone bead. DNA-bending non-histone protein is modeled as a spring (red) connecting two beads in the linker region.

The DNA polymer is modeled as a bead-spring chain having *N*_*d*_ beads connected by *N*_*d*_ - 1 springs (Fig. 1). A nucleosome is modelled as one big bead representing the core histone octamer on which 14 DNA beads are wrapped around in 1.75 turns. To explicitly model histone tails, we introduce eight small flexible polymers emanating from the core histone as shown in Fig. 1. The DNA polymer bead has a diameter of 3.4 nm while the octamer core histone bead has a diameter of 5.25 nm. The details of each histone-tail polymer length is as follows: H2A tail = 6.2 nm (4 beads), H2B tail = 7.8 nm (5 beads), H3 tail = 12.6 nm (8 beads), and H4 tail = 7.8 nm (5 beads) (21). We fix the first bead of each histone tail on the core histone bead such that the first bead and the core histone bead behave like a rigid body. We introduce the linker histone H1 as a separate bead (diameter = 2.9nm (44)), which interacts via a harmonic spring with the two DNA beads entering and exiting the nucleosome as well as with the nucleosome. The H1 bead constrains the DNA tangent vectors entering and exiting the nucleosomes ensuring known structural features of nucleosomes.

The DNA polymer is modeled as a bead-spring chain having *N*_*d*_ beads connected by *N*_*d*_ - 1 springs (Fig. 1). A nucleosome is modelled as one big bead representing the core histone octamer on which 14 DNA beads are wrapped around in 1.75 turns. To explicitly model histone tails, we introduce eight small flexible polymers emanating from the core histone as shown in Fig. 1. The DNA polymer bead has a diameter of 3.4 nm while the octamer core histone bead has a diameter of 5.25 nm. The details of each histone-tail polymer length is as follows: H2A tail = 6.2 nm (4 beads), H2B tail = 7.8 nm (5 beads), H3 tail = 12.6 nm (8 beads), and H4 tail = 7.8 nm (5 beads) (21). We fix the first bead of each histone tail on the core histone bead such that the first bead and the core histone bead behave like a rigid body. We introduce the linker histone H1 as a separate bead (diameter = 2.9nm (44)), which interacts via a harmonic spring with the two DNA beads entering and exiting the nucleosome as well as with the nucleosome. The H1 bead constrains the DNA tangent vectors entering and exiting the nucleosomes ensuring known structural features of nucleosomes.

Now we describe the energetics of the chromatin system that we consider. Throughout this paper, we will use notation with subscript (*α, β*) ∈ {*d, h, t, l*} where *d, h, t, l* stand for the DNA bead, histone bead, tail bead, and linker bead, respectively. For example, the diameter of a DNA bead will be represented by *σ*_*d*_ while that of the tail bead will be *σ*_*t*_, and so on. All beads in the system are connected with their respective neighbours using a harmonic potential given by the general formula as:

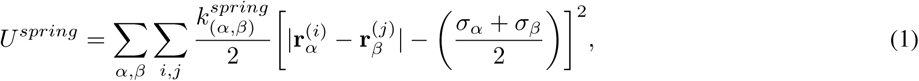

where 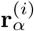 and 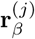 are position vectors of *i*^th^ and *j*^th^ beads of types *α* and 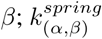 is the corresponding spring constant. (i) For DNA–DNA bead interaction *α* = *d, β* = *d, j* = *i* + 1 and *i* ranges from 1 to *N*_*d*_ - 1. (ii) For histone core-DNA interaction *α* = *d, β* = *h, i* ranges from 1 to *N*_*h*_ (where *N*_*h*_ is total number of nucleosomes) and *j* takes 14 values, for each *i*, representing the identity of the corresponding histone-bound DNA beads. (iii) For tail–tail bead interaction *α* = *t, β* = *t, j* = *i* + 1 and *i* = *N*_*t*_ - 1 where *N*_*t*_ is number of the tail bead. (iv) For DNA and linker histone interaction *α* = *l, β* = *d, i* ranges from 1 to *N*_*l*_ (where *N*_*l*_ is total number of linker histones), *j* stands for 3 DNA bead positions at the entry site, exit site, and the center (where the DNA bead is wrapped around the histone) along the dyad axis. The interaction energy of the NHPs is also calculated using equation 2. A NHP binds at any linker region between two linker DNA beads *α* and *β*. If there are *ν* NHPs, each having a size spanning 3 beads, *i* is the count of the NHPs varying from 1 to *ν* and *j* = *i* + 3, and *v/N*_*h*_ is called protein density. The total spring energy (*U*^*spring*^) is obtained by summing over with appropriate nearest neighbour bead pairs *α, β*. Apart from the nearest neighbour spring interaction described above, any two beads interact via the standard Lennard-Jones (LJ) potential (*U*^*LJ*^) such that 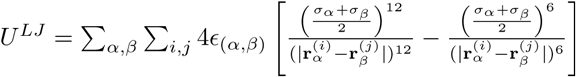, when 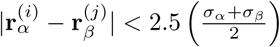, and *U*^*LJ*^ = 0 otherwise. That is, the energy is zero when 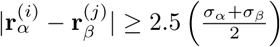. Here *ϵ*_(*α,β*)_is the corresponding potential well depth. All beads interact with each other using screened electrostatic potential. We use the standard Debye–Hückel potential and compute this energy of chromatin (21) as follows: 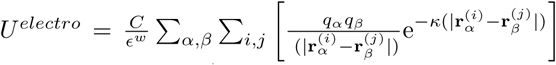, where *q*_*α*_ and *q*_*β*_ are effective charges of beads, *κ* is the inverse of the debye length (1nm^*-*1^), *ϵ*^*w*^ is the dielectric constant (set to 80 assuming a water-like medium) and C is a constant as per the screened Coulomb electrostatic potential energy formula (45). Total charges on wrapped DNA are estimated as *-* 296*ϵ* (46). 14 DNA beads wrap around the core histone, so charge value for one DNA bead *q*_*d*_ = − 296*/*14 = −21.14*e*. The core histone bead (without tails) has a charge of *q*_*h*_ = 52*e* (46). We take charge on each histone-tail bead *q*_*t*_ = 2*e* such that the total charge on one nucleosome (wrap DNA+core histone+histone tails) is maintained as 156 (46). Charges for histone H1 are taken as *q*_*l*_ = 13.88 (21).

The semiflexible nature of the DNA chain is introduced through a bending potential (*U*^*bend*^) for DNA beads as: 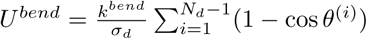 where *k*^*bend*^ is the bending stiffness of DNA and *θ*^(*i*)^ is the angle between two nearby bonds in the bead-spring model.

The total energy of the chromatin in this model is given by *U*^*tot*^ = *U*^*spring*^ + *U*^*electro*^ + *U*^*LJ*^ + *U*^*bend*^. This system was simulated using Molecular Dynamics simulation package LAMMPS (47) and we obtained chromatin configurations (3D positions of all beads) as a function of time. Most of the parameters used in simulations are taken from known experimental data or earlier computational studies. Apart from the bead sizes and charges mentioned earlier, the spring constants are assumed to have a large value 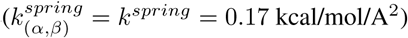 such that the bonds are stable and does not fluctuate a lot. The time step of the simulation is Δ*t* = 359.5*fs*. The details about the parameters are given in the *SI Appendix*, Table 1.

Now we describe the energetics of the chromatin system that we consider. Throughout this paper, we will use notation with subscript (*α, β*) ∈ {*d, h, t, l*} where *d, h, t, l* stand for the DNA bead, histone bead, tail bead, and linker bead, respectively. For example, the diameter of a DNA bead will be represented by *σ*_*d*_ while that of the tail bead will be *σ*_*t*_, and so on. All beads in the system are connected with their respective neighbours using a harmonic potential given by the general formula as:

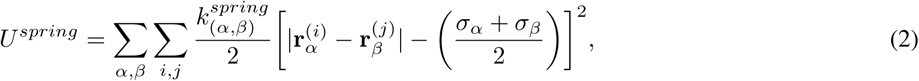

where 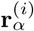 and 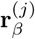 are position vectors of *i*^th^ and *j*^th^ beads of types *α* and 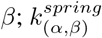 is the corresponding spring constant. (i) For DNA–DNA bead interaction *α* = *d, β* = *d, j* = *i* + 1 and *i* ranges from 1 to *N*_*d*_ - 1. (ii) For histone core-DNA interaction *α* = *d, β* = *h, i* ranges from 1 to *N*_*h*_ (where *N*_*h*_ is total number of nucleosomes) and *j* takes 14 values, for each *i*, representing the identity of the corresponding histone-bound DNA beads. (iii) For tail–tail bead interaction *α* = *t, β* = *t, j* = *i* + 1 and *i* = *N*_*t*_ - 1 where *N*_*t*_ is number of the tail bead. (iv) For DNA and linker histone interaction *α* = *l, β* = *d, i* ranges from 1 to *N*_*l*_ (where *N*_*l*_ is total number of linker histones), *j* stands for 3 DNA bead positions at the entry site, exit site, and the center (where the DNA bead is wrapped around the histone) along the dyad axis. The interaction energy of the NHPs is also calculated using equation 2. A NHP binds at any linker region between two linker DNA beads *α* and *β*. If there are *v* NHPs, each having a size spanning 3 beads, *i* is the count of the NHPs varying from 1 to *v* and *j* = *i* + 3, and *v/N*_*h*_ is called protein density. The total spring energy (*U*^*spring*^) is obtained by summing over with appropriate nearest neighbour bead pairs *α, β*. Apart from the nearest neighbour spring interaction described above, any two beads interact via the standard Lennard-Jones (LJ) potential (*U*^*LJ*^) such that 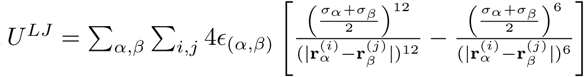, when 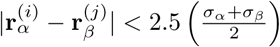, and *U*^*LJ*^ = 0 otherwise. That is, the energy is zero when 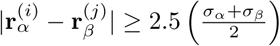. Here *ϵ*_(*α,β*)_ is the corresponding potential well depth. All beads interact with each other using screened electrostatic potential. We use the standard Debye–Hückel potential and compute this energy of chromatin (21) as follows: 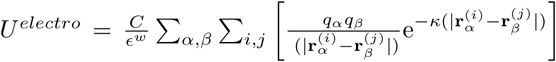, where *q*_*α*_ and *q*_*β*_ are effective charges of beads, *κ* is the inverse of the debye length (1nm^*-*1^), *ϵ*^*w*^ is the dielectric constant (set to 80 assuming a water-like medium) and C is a constant as per the screened Coulomb electrostatic potential energy formula (45). Total charges on wrapped DNA are estimated as *-* 296*ϵ* (46). 14 DNA beads wrap around the core histone, so charge value for one DNA bead *q*_*d*_ = *-* 296*/*14 = *-* 21.14*ϵ*. The core histone bead (without tails) has a charge of *q*_*h*_ = 52*e* (46). We take charge on each histone-tail bead *q*_*t*_ = 2*ϵ* such that the total charge on one nucleosome (wrap DNA+core histone+histone tails) is maintained as 156*ϵ* (46). Charges for histone H1 are taken as *q*_*l*_ = 13.88 (21).

The semiflexible nature of the DNA chain is introduced through a bending potential (*U*^*bϵnd*^) for DNA beads as: 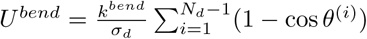 where *k*^*bϵnd*^ is the bending stifness of DNA and *θ*^(*i*)^ is the angle between two nearby bonds in the bead-spring model.

The total energy of the chromatin in this model is given by *U*^*tot*^ = *U*^*spring*^ + *U*^*ϵlϵctro*^ + *U*^*LJ*^ + *U*^*bϵnd*^. This system was simulated using Molecular Dynamics simulation package LAMMPS (47) and we obtained chromatin configurations (3D positions of all beads) as a function of time. Most of the parameters used in simulations are taken from known experimental data or earlier computational studies. Apart from the bead sizes and charges mentioned earlier, the spring constants are assumed to have a large value 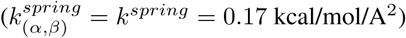 such that the bonds are stable and does not fluctuate a lot. The time step of the simulation is Δ*t* = 359.5*fs*. The details about the parameters are given in the *SI Appendix*, Table 1.

## Results

We present below the results from our study, using computer simulations, where we examined how the chromatin organization alters as we vary different interaction potential energies (electrostatic, LJ), DNA-bending due to NHPs, and the length of the interacting domain. We will discuss how heterogeneity in interaction potential energies will lead to heterogeneous chromatin configurations.

### Role of different interactions in chromatin packaging

To mimic interactions due to histone modifications, presence of various proteins, and many other factors, we simulated the system introducing electrostatic interaction energy and Lennard-Jones interaction energy into our model.

First we present results from a simulation of a chromatin made of 20 nucleosomes, accounting for histone tails, and H1 linker histones, but not considering the DNA-bending NHPs. 3D configurations of such a chromatin for different values of interaction potential strengths are shown in Fig. 2(a). In the left panel within Fig. 2(a), we have chromatin for different LJ interaction strengths but with no electrostatic interactions. In the right panel, chromatin configurations for exactly the same condition but with electrostatic interactions are shown. When electrostatic interaction is present, the chromatin is more compact. The positive charges of the histone tails attract with negatively charged parts of the chromatin. This leads to a compact structure.

**Figure 2.**
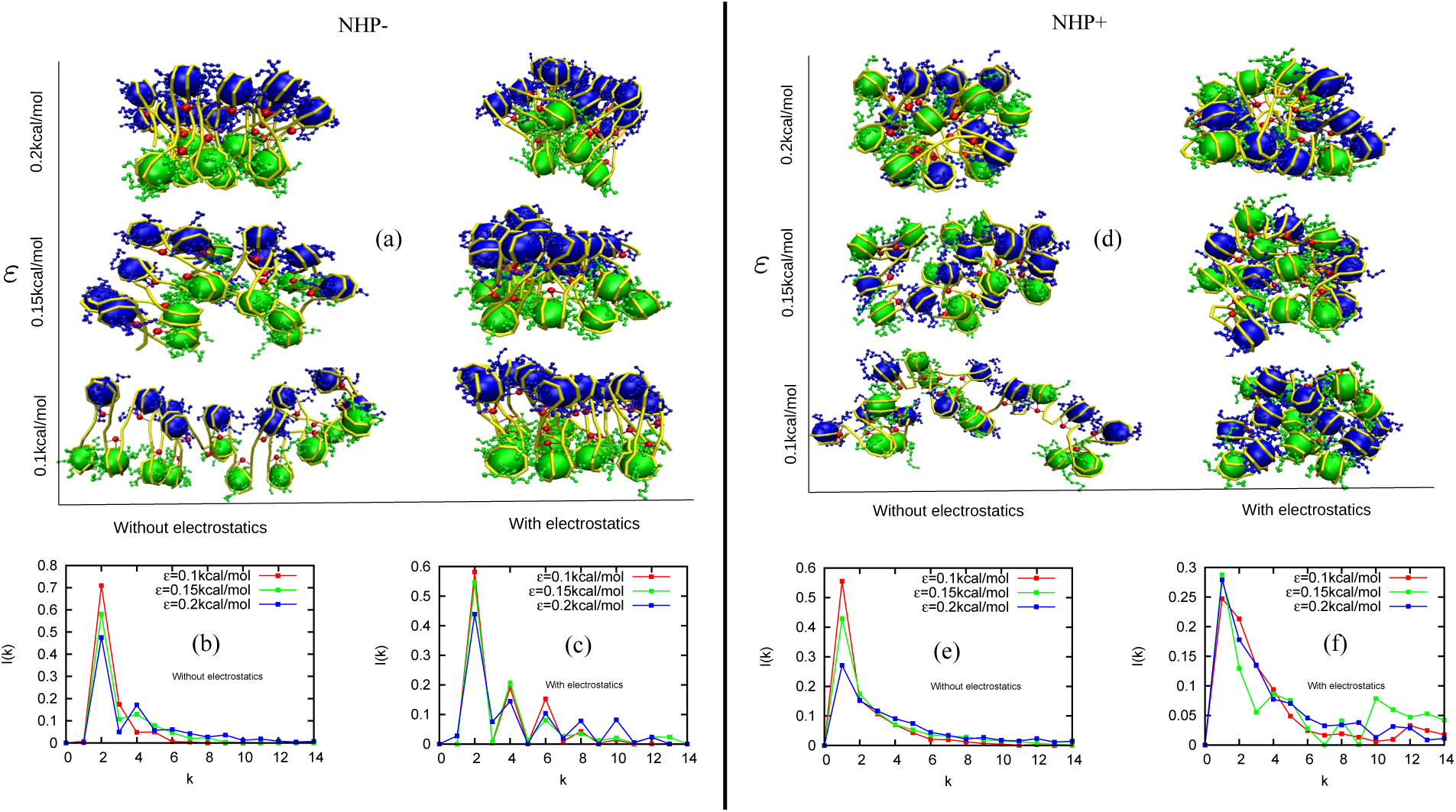
Snapshots of chromatin configurations and the corresponding contact probabilities between *k*^*th*^ neighbour nucleosomes (*I*(*k*)) in the absence (NHP-) and presence (NHP+) of DNA-bending non-histone proteins (NHPs) that bind along linker DNA regions. The results are presented for four cases: case 1: Simulations with no NHPs and no electrostatic interactions (left column of (a), and (b)); case 2: No NHPs but with electrostatic interactions (right column of (a), and (c)); case 3: with NHPs and no electrostatic interactions (left column of (d), and (e)); case 4: With NHPs and with electrostatic interactions (right column of (d), and (f)). All the results are presented for 3 different strengths of LJ (*ϵ*) interaction potentials (see text for details).

Based on nucleosome structure data, we know that each tail-bead in our coarse-grained model, has one or more sites that can be acetylated. Acetylation will reduce electrostatic interaction in the tails. Our “without electrostatics” results are comparable to configurations of a chromatin with highly acetylated nucleosomes; we get open structures when there is no electrostatic interactions and relatively more compact structures with electrostatic interactions analogous to chromatin with acetylated and de-acetylated nucleosomes. This is consistent with earlier findings (5, 42).

If we go along the column from bottom to top (Fig. 2(a)), we find chromatin organization for different values of *ϵ* (LJ interaction strength). For small values of *ϵ*, chromatin is more open while for larger values of LJ interaction strengths, the chromatin is compact. Here we introduced attractive part of the LJ potential to mimic different types of interactions mediated by various proteins (like HP1) or other interactions that may bring together nucleosomes (48). The repulsive part of LJ mimics steric hindrance.

We then computed *I*(*k*)—the probability that a nucleosome is in contact with its *k*^th^ neighbour— from the above-discussed 3D chromatin configurations (see *SI Appendix* and Figs. 2(b) & (c)). We find that, irrespective of electrostatic interactions and the strength of the LJ interactions, the most prominent peak is at *k* = 2. Note that when we change the strength of the LJ, in the absence of electrostatics, the height of the prominent peak varies. This is indeed related to the compact packaging of the zig-zag structure. In the open form (low LJ), the zig-zag is more loose; hence, only next-neighbor nucleosomes (k=2) mainly interact; far away nucleosomes (*k >* 2) are not contributing to *I*(*k*). However, for large LJ, the chromatin is compact and far away nucleosomes have a higher chance of being close and hence, *I*(*k*) values for *k >* 2 are larger. Since total probability is 1(Σ_*k*_ *I*(*k*) = 1), this reduces the peak value of I(2) for large LJ values. Similarly, when electrostatic interactions are present, the compactness of the chromatin is reflected in higher *k >* 2 peaks.

### Even compact chromatin structures are irregular in the presence of DNA-bending NHPs

Here we investigate the interplay between DNA-bending due to NHPs and electrostatic/LJ interactions in deciding the 3D folding of chromatin. We did simulations similar to the ones described earlier, but introduced NHPs that bind in the linker region and bend DNA, assuming NHP density of 0.5 per linker region. Typical snapshots of chromatin 3D configurations are shown in Fig. 2(d) with (right panel) and without (left panel) electrostatic interactions; also see the LJ variation along the vertical axis. Without electrostatics, for small LJ, the chromatin is in open configuration. As we increase the LJ interaction (high *ϵ*), the chromatin becomes more compact. When electrostatic interactions were switched on (right panel), we got relatively compact structures for all values of *ϵ*.

The important point to note is that in this figure (Fig. 2(d)), as opposed to the earlier case in Fig. 2(a), different color beads (odd and even numbered beads with blue and green colors, respectively) are well mixed, suggesting a irregular organization of chromatin. The presence of NHPs have created an irregular chromatin even with electrostatic interactions and high LJ interactions.

Quantification of the 3D structure using contact probability *I*(*k*) shows that, in the presence of NHPs, irrespective of electrostatic interactions and the strength of the LJ interactions, the most prominent peak is at *k* = 1 (Fig. 2(e)). It is interesting to see that even for the high compact form of chromatin, structures are irregular in the presence of NHPs. Note that when we change the strength of the LJ, with no electrostatic interaction, the height of the prominent peak (peak at *k* = 1) varies. This is indeed related to the compact packaging of the irregular structure. In the open form (low LJ), the irregular structure is more loose, hence only neighbor nucleosomes (k=1) mainly interact; far away nucleosomes (*k >* 2) are contributing relatively less to *I*(*k*). However, for large LJ, the chromatin is compact and far away nucleosomes have a higher chance of being close and hence, *I*(*k*) values for *k >* 1 are higher. Since total probability is 1 (Σ_*k*_ *I*(*k*) = 1), this reduces the peak value of *I*(1) for large LJ values. In Fig. 2(f), with electrostatic interactions, the structure is highly compact and irregular; hence the *I*(*k*) peaks are roughly the same for all values of LJ interaction strengths.

### Regular vs. irregular chromatin: packing density, fiber width, and mean cluster size

We computed packing density to know compaction of different chromatin structures for the discussed earlier cases. Packing density measures roughly the number of nucleosomes packed in every 11nm of effective chromatin length (29). First we consider chromatin organization with no NHPs; these are regular ordered chromatin structures as seen in the earlier results (Fig. 2(a)). In the absence of NHPs and in the presence of electrostatic interactions, the packing density is *≈* 6 nucleosomes/11nm (see Fig. 3(a), green curve) as expected for zig-zag or regular chromatin (11, 16, 17, 23–27). The results show that the chromatin is highly packed even for a very small LJ interaction strength(*ϵ* = 0.1kcal/mol) and remains nearly the same for larger LJ interaction strengths (*ϵ* = 0.2kcal/mol). In the absence of electrostatic interactions (Fig. 3(a), red curve), packing density is much smaller (*≈* 2 *-* 3 nucleosomes/11nm) than the case with electrostatic interactions suggesting that the electrostatics plays an important role in packaging. On increasing the strength of the LJ interactions from *ϵ* = 0.1kcal/mol to *ϵ* = 0.2kcal/mol, we found that the packing density increased slightly, but remained smaller than that of the chromatin with electrostatic interactions.

**Figure 3.**
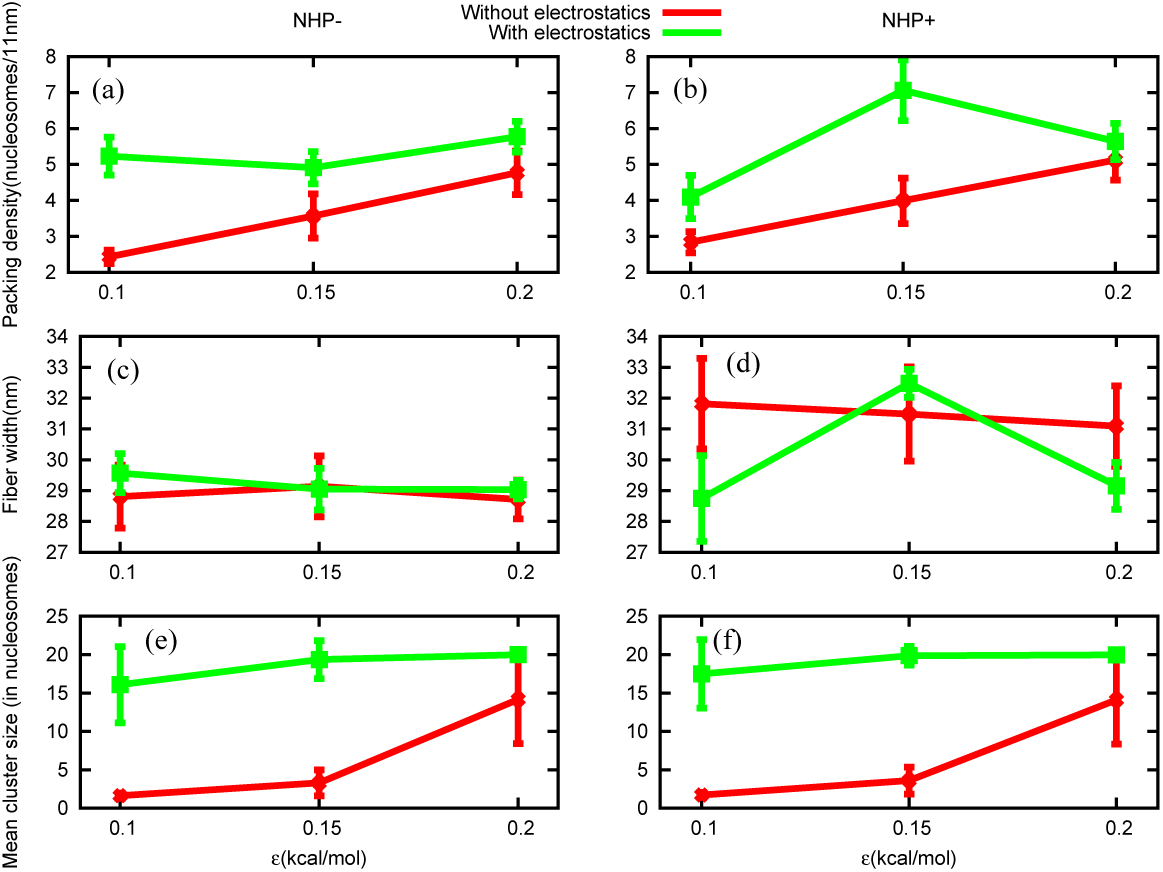
Packing density, fiber width, and mean cluster size are plotted for different LJ strengths (*ϵ*), in the absence of NHPs (NHP-, left side) and presence of NHPs (NHP+, right side). (a),(b) Without electrostatic interaction, the packing density increases with *ϵ*(red curves). With electrostatic interaction, packing density is *≈* 6 nucleosomes/11nm (green curve). (c),(d) Chromatin fiber diameter (width) for different parameters. With (green curve) and without (red curve) electrostatic interaction, the fiber diameter is *≈* 30 nm (constant) for all cases. (e),(f) Without electrostatic interaction, mean cluster size increases on increasing the LJ (*ϵ*) parameter (red curve), but with electrostatic interaction it remains constant(green curve). In all the subfigures, the vertical bars represent standard deviation.

With DNA-bending NHPs, we got a packing density of 4 to 7 nucleosomes/11 nm, in the presence of electrostatic interactions (Fig. 3(b)). This implies that, even though our models with NHPs do not give rise to regular/ordered structure, the packing/compaction remains similar to that of what is observed in *in vitro* experiments. Our theory predicts that DNA-bending NHPs will make chromatin irregular but will have similar compaction as observed in the case without NHPs (11, 16, 17, 23–27). The findings for the case with no electrostatic interactions are also similar to the case without NHPs —a lower packing density that increases as a function of LJ interaction strengths.

In Fig. 3(c) and (d), we present our results for fiber width in the absence and presence of NHPs, with (green curve) and without (red curve) electrostatics. Interestingly, for all the cases, the width is *≈* 30nm. This width is computed according to a simple definition based on polymer physics ideas (see *SI Appendix* for details). We know (from Fig. 2(b)) that in the presence of NHPs, chromatin is irregular; we also know that in the absence of electrostatic interactions, the chromatin is relatively more open. However, in all these cases the width is approximately 30nm. This shows that the width as a quantity, as defined here, cannot easily distinguish between regular and irregular, open and compact chromatin. However, note that the packing density in the Fig. 3(a) and (b) could distinguish between open and compact chromatins (red and green curves).

Since both compact and open structures are seen to have similar width, to improve the quantification, we computed how nucleosomes are clustered near one another. Any two nucleosomes that are closer than 2.5 times its diameter (2.5*σ*_*h*_) are considered to be in the same cluster (see *SI Appendix*). We find that the open chromatin is not just one cluster but many small clusters having a few nucleosomes in each—see Figs. 3(e) & (f) (red curve) where the mean cluster size is small (*≈* 2 *-* 5) and hence, the mean cluster number is large (4 *-* 10). On the other hand, the compact chromatin is nearly one cluster — *≈* 20 nucleosomes are part of the same cluster (see Fig. 3(e),(f), green curve). This suggests that open chromatin, even through it appears like a loose zig-zag visually, may not appear as a single entity (punctate) in experiments (like cryo-EM), where the density and cluster size can affect the measurement.

### Role of chromatin length in packing density and fiber width

So far, we studied packing density, fiber width, and cluster size for a chromatin having 20 nucleosomes. We found that, even though the NHPs make the chromatin irregular, a compact chromatin fiber has a packing density *≈* 6 nucleosomes/11 nm, and a width of *≈* 30 nm. How do packing density, width of the fiber, and other parameters vary as we change the length of the polymer? In this section we simulated chromatin fibers of many different lengths starting with chromatin having 12 nucleosomes going upto 100 nucleosomes. In Fig. 4(a), snapshots of the chromatin structure for polymers having 50 nucleosomes and 100 nucleosomes are shown in the absence and presence of NHPs. Packing densities and fiber widths for different lengths (12, 20, 50, and 100 nucleosomes) are shown in Fig. 4(b). In the absence (left panel) and presence (right panel) of NHPs, packing densities and fiber widths increase on increasing the polymer length from 12 nucleosomes to 100 nucleosomes. The longer the polymer, the more the width and higher the packing density. For polymers having 20 or more nucleosomes, the width is 30 nm or above; the packing density is above 6 nucleosomes per 11 nm. This result suggests that if we pack chromatin, even in an irregular manner, with high compaction, one will get structures having width 30 nm or more. However, in the context of *in vivo* chromatin, where recent experiments do not find chromatin of width 30nm or more, this presents a puzzle. Why is it that in *in vivo* experiments we do not find chromatin a having width 30 nm? Recent *in vivo* experiments have found chromatin of widths only in the range 5–24 nm (37).

**Figure 4.**
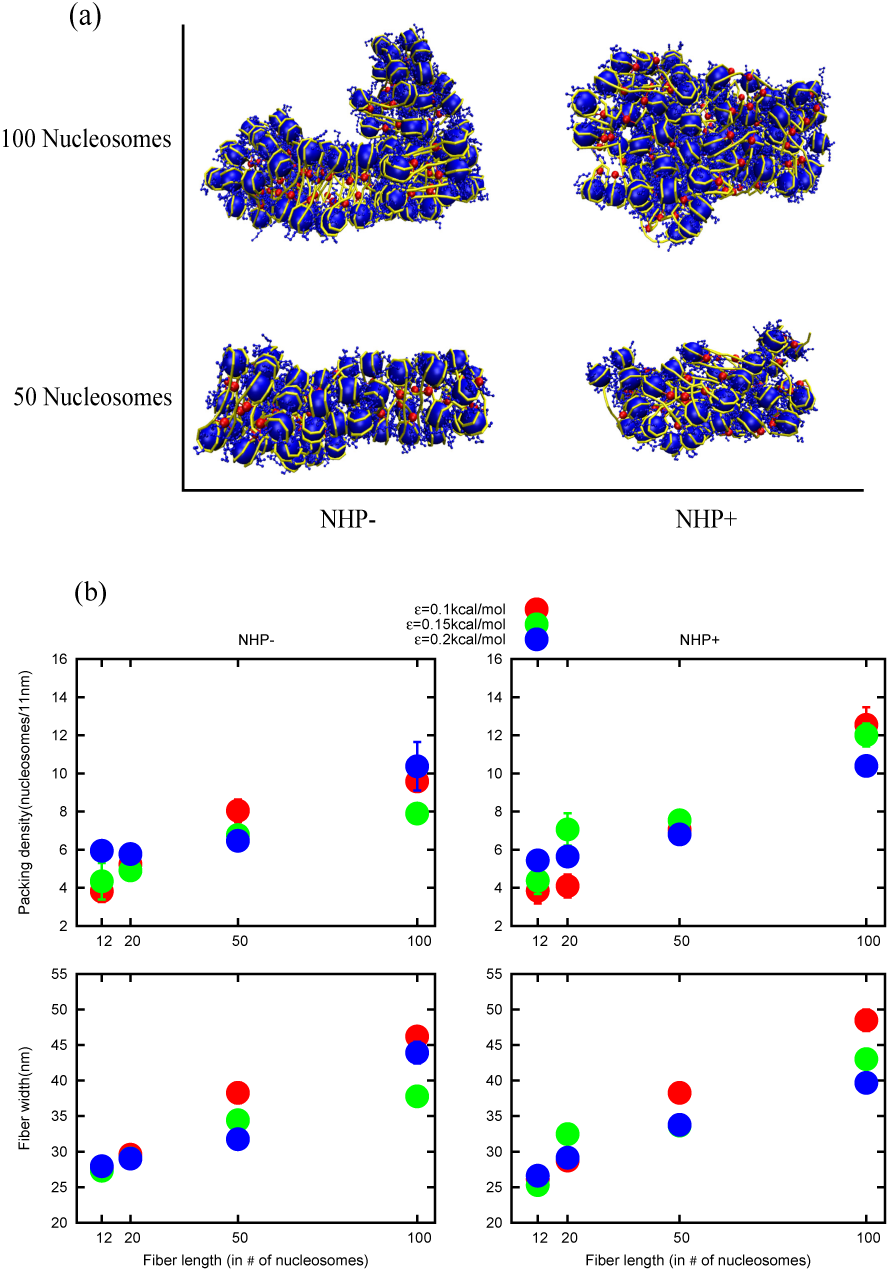
(a) Snapshots of simulation results of 50 nuclesomes and 100 nucleosomes in the absence and presence of NHPs (left to right). (b) Packing density and fiber width on varying chromatin fiber length in the absence and presence of NHPs for different LJ interaction strengths (*ϵ*). Packing densities and fiber widths increase as we increase the chromatin length.

### Spatial variation in histone modifications and interaction potentials

To address the puzzle mentioned earlier, where one gets 30 nm wide chromatin if we have highly interacting 20 or more nucleosomes, we turn our attention to the histone modifications. We know that histone modifications vary spatially (along the contour of a chromatin) and can influence the nature of effective interactions between the different parts of the chromatin (49). The presence of certain methylations (like H3K9me3) could lead to recruitment of certain proteins (like HP1) and induce high attractive interactions between chromatin segments (48). The absence/presence of acetylations could also affect the local electrostatic potentials leading to heterogeneous interactions. All these will affect the packaging and clustering of nucleosomes. To understand this heterogeneity in interactions, as an example, we examined the extent of H3K9me3 modifications along the chromatin fiber length. Using ChIP-seq data available in public databases, we asked the following question. What is the length of the typical contiguous patch of chromatin having H3K9me3 modification?(human T-cell in ref. (50)). We defined any two modification peaks as a part of the same contiguous patch if the separation between the peaks is less than 1000 bp. Using this definition, we computed length distribution of contiguous patches of chromatin having H3K9me3 modifications for 22 human chromosomes (see Fig. 5(a)). The distributions peak at small values of lengths suggesting that very long contiguous patches are rare. The mean length of H3K9me3 modification patch across all chromosomes is *≈* 1600 bp and the gap length between two contiguous patches is *≈* 2200 bp. If we convert in terms of nucleosomes, this is equivalent to chromatin having *≈* 8 nucleosomes with modifications and *≈* 12 nucleosomes without.

**Figure 5.**
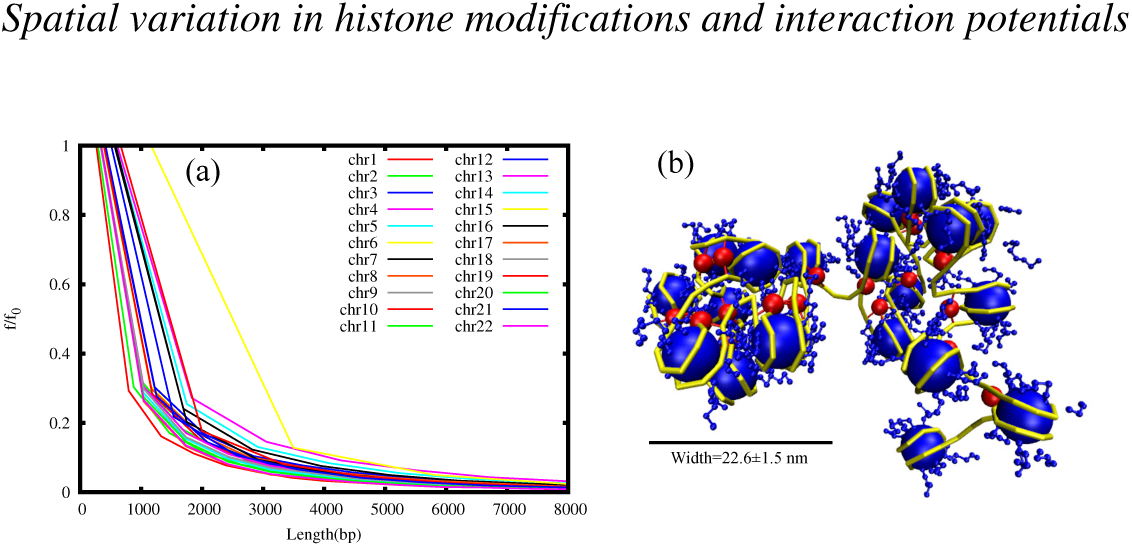
(a) Frequency (*f*) distribution of lengths of contiguous patches having H3K9me3 modifications, for different chromosomes, scaled with maximum frequency (*f*_0_). Most of the patches are smaller in length; very long patches are rare. (b) A snapshot of the chromatin structure with NHPs and heterogenous interactions: the system was simulated with electrostatic interactions in the first 8 nucleosomes, and with no electrostatic interaction in the remaining 12 nucleosomes.

To mimic the essence of spatial variation in modifications, we simulated 20 nucleosomes where we introduced electrostatic interactions into one fraction (8 nucleosomes) and the other fraction (12 nucleosomes) was left with no electrostatic interactions. The LJ potential was kept uniform (*ϵ* = 0.1kcal/mol) and the linker regions were bent with NHPs having a density of 0.5. The resulting chromatin structure is shown in Fig. 5(b). The 8 nucleosomes having electrostatic interactions form one cluster, while the other 12 nucleosomes divided into many small clusters. Note that here the chromatin is irregular due to the presence of NHPs. We calculated the width of the single cluster of 8 nucleosomes which is 22.6 *±* 1.5 nm. The other small clusters have widths varying from a single-nucleosome width (*≈* 5–10 nm) to two or three nucleosome widths (10–20 nm). This results are comparable to recent experimental results where they found predominantly fiber width to be in the range of 5 *-* 24 nm (37). The distribution of the contignous patch lengths of histone modifications suggest that chromatin configurations must be computed with non-uniform interaction potentials and it will result in a chromatin where most of the regions have widths *<* 30 nm; regions of higher width will be rare. In other words, heterogeneity in interactions resulting from actively maintained spatial variation of histone modifications may determine the width of chromatin fibers *in vivo*.

## Conclusion

In this work we examined the width, packing density, and clustering properties of chromatin that is irregular due to the binding of DNA-bending NHPs. We find that the DNA-bending NHPs will make chromatin irregular; however, the resulting structure can have similar packing density and width as that of the chromatin without NHP (regular). We examined how different factors such as the length of the chromatin and the nature/strength of interactions determine the width, packing density, and clustering properties of chromatin. In our simulations, we explicitly accounted for histone tails, and electrostatic interaction and varied the length of chromatin across several nucleosomes (length of a typical gene to many genes). We showed that electrostatic interactions make the chromatin more compact, whereas a chromatin simulated without electrostatic interactions is more open. This is consistent with the fact that reduction in net positive charges (hence reduction in electrostatic interactions) due to acetylation of histone tails leads to more open chromatins. We examined how the nucleosomes are clustered in a highly packed chromatin and in an open chromatin. We calculated the mean cluster size of simulated structures and showed that open chromatin is essentially many small clusters of nucleosomes; they may not appear as a single thick fiber in many experiments. We varied fiber length and calculated packing densities and fiber widths and showed that both quantities increase with fiber length. Then we addressed the resulting puzzle as to why one does not observe highly packed chromatin fibers of width 30 nm or above *in vivo*? We argued that one of the missing components here could be the heterogeneity in interactions resulting from histone modifications. We simulated chromatin configurations considering this heterogeneity in interactions and showed that heterochromatin structure could have a typical width less than 30 nm.

The other factor that might affect the local structure and the width is the ATP-dependent activity and dynamics of the nucleosomes and proteins. In this work, we have assumed that nucleosomes and NHPs are static, and do not change their organization with time. However, remodeling machines and cellular processes like gene-expression can disassemble/ reorganize nucleosomes, which may affect the properties of chromatin in the lengthscale of a gene. However, our model will be a reasonable description for inactive regions—regions where there are least remodeling activity and gene expression. It is believed that such heterochromatic regions will have 30 nm wide chromatin structures. However, our work suggests that even there, depending on the nature of histone modifications, one may get chromatin widths smaller than 30 nm. We have assumed that electrostatic interactions obey a Debye–Hückel potential; however, fluctuations of counter-ions and other charged constituents are not accounted for in this description of the interactions. These are some of the limitations of our model.

### Suggestion for new experiments to test our prediction

Experimentally, our findings can be tested in a few different ways. One can possibly perform *in vitro* chromatin reconstitution experiments with NHPs and measure the width, packing density, and cluster sizes. Our prediction is that the chromatin will be irregular, but with a width of *≈* 30 nm, if the length is appropriately chosen as we have shown in our results; however depending on the nature of the modifications, one may not find the chromatin as one cluster but many small clusters of nucleosomes. This may be repeated for many different lengths and quantities measured as a function of length. Our work also suggests that one should experiment with heterogeneity in interactions; this may be introduced by appropriately mutating charged/neutral amino acids in the tail region in a fraction of the histones. This will bring heterogeneity in electrostatic interactions, and according to our predictions this can lead to alterations in the width and packing density of chromatin.

To conclude, in this work, we simulated chromatin in the lengthscale of a few genes, accounting for various factors. Our results show the importance, heterogeneity in interactions and the role of NHPs. This work should be considered as a step in the direction toward a more complete model to study the chromatin states and the dynamics of chromatin accounting for realistic details like protein-binding and interactions due to histone modifications. We hope that this work will lead to further experimentation and computation.

## Supporting information

