## Supplementary material for "Irregular chromatin: packing density, fiber width and occurrence of heterogeneous clusters"

### 1 Contact probability $I(k)$

We define $I(k)$ which is the probability that any nucleosome is in "contact" with its $k^{th}$ neighbor. More precisely $I(k)$ is the probability of finding $k^{th}$ neighbor nucleosome below a certain cut-off distance (here the cut off distance is taken as $2.5\sigma_h = 13.2$nm). To compute this probability, (similar to the procedure followed in ref [1]), we first define a square matrix $D^{i,j}$ which has elements 1 or 0 and is defined by:

$$D^{i,j}(n) = \begin{cases} 1 & \text{if } |\mathbf{r}_h^{(i)} - \mathbf{r}_h^{(j)}| < 2.5\sigma_h \\ 0 & \text{else} \end{cases} \tag{1}$$

where $n$ is the $n^{th}$ configuration (obtained from simulations), and $\mathbf{r}_h^{(i)}$ and $\mathbf{r}_h^{(j)}$ are the positions of $i^{th}$ and $j^{th}$ nucleosome. Let $\overline{D}(i,j)$ be the average of this matrix over different configurations ($n$), in stead-state. Then we compute I(k) which is nothing but the probability of $k^{th}$ neighbor nucleosome below a cut-off distance of $2.5\sigma_h$, as:

$$I(k) = \frac{\tilde{I}(k)}{\sum\limits_j \tilde{I}(j)} \tag{2}$$

where $\tilde{I}(k) = \sum\limits_{i=1}^{N_h-k} \overline{D}(i, i+k)$ and $N_h$ is total number of nucleosomes present in the chromatin.

### 2 Packing density and fiber width

We compute packing density $p_d$, which is roughly defined as the number of nucleosomes ($N_h$) packed in every 11nm effective length ($L_{fiber}$) of the chromatin fiber.

$$p_d = \frac{N_h \times 11\text{nm}}{L_{fiber}} \text{nucleosomes/11nm}, \tag{3}$$

To calculate effective length ($L_{fiber}$) of the chromatin fiber, (similar to the procedure followed in ref [1]), we define the fiber axis $\mathbf{r}_{ax} \approx (P_x^{(i)}, P_y^{(i)}, P_z^{(i)})$. $\mathbf{r}_{ax}$ is calculated by solving equations of polynomial $P_x^{(i)}, P_y^{(i)}$ and $P_z^{(i)}$ from least square method which is best fit with $i^{th}$ nucleosome position $\mathbf{r}_h^{(i)} = (x_h^{(i)}, y_h^{(i)}, z_h^{(i)}$ in the x, y and z direction respectively (see Fig. S1). Then we computed fiber length $L_{fiber}$ as follows:

$$L_{fiber} = \sum_{i=1}^{(N_h-1)/2} |\mathbf{r}_{ax}^{(2i-1)} - \mathbf{r}_{ax}^{(2i+1)}|, \tag{4}$$

We also calculate fiber width ($w_d$) which is Following:

$$w_d = \frac{2}{N_h} \sum_{i=1}^{N_h} |\mathbf{r}_h^{(i)} - \mathbf{r}_{ax}^{(i)}| + 5.5\text{nm}. \tag{5}$$

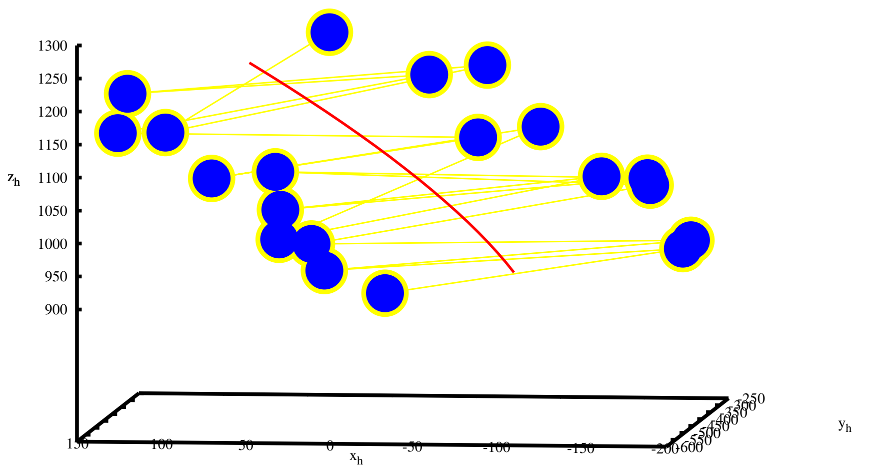


(a)

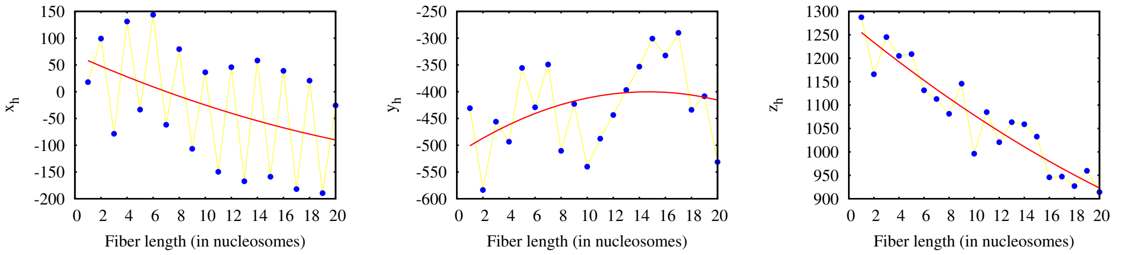


(b) (c) (d)

Fig. S1: (a) Fiber axis (red curve) is calculated using least square method which is best fitted with nucleosome center positions (blue dots). Yellow lines are connections between two consecutive nucleosomest. (b),(c) and (d) Fiber axis decomposition (red curve) with center position of nucleosomes (blue dots) into x, y and z directions respectively.

### 3 Cluster of nucleosomes

When a few nucleosomes come within a certain cutoff distance (we took 2.5 times its diameter, $2.5\sigma_h$), they are defined as a cluster of nucleosomes. If a nucleosome has no neighbor within the cutoff distance, it is considered to be a cluster of size 1. Similarly if a nucleosome has $N_h - 1$ neighbors within the cutoff distance, it is considered as a cluster size $N_h$.

### 4 Simulation parameter description

| **Parameter** | **Description** | **Value** |
|---|---|---|
| $q_d$ | Charge on DNA bead | $-21.14e$ [1] |
| $q_h$ | Charge on core-histone bead | $52e$ [2] |
| $q_t$ | Charge on histone tail bead | $2e$ |
| $q_l$ | Charge on linker histone(H1) bead | $13.88e$ [1] |
| $\sigma_d$ | Diameter of DNA bead | $34\text{A}^o$ [1] |
| $\sigma_h$ | Diameter of core-histone bead | $52.5\text{A}^o$ [3] |
| $\sigma_t$ | Diameter of histone tail bead | $15.6\text{A}^o$ [1] |
| $\sigma_l$ | Diameter of linker histone(H1) bead | $29\text{A}^o$ [4] |
| $m_d$ | Mass of DNA bead | 6000gm/mole [3] |
| $m_h$ | Mass of core-histone bead | 22089gm/mole |
| $m_t$ | Mass of histone tail bead | 579gm/mole |
| $m_l$ | Mass of linker histone(H1) bead | 5118gm/mole |
| $k^{spring}$ | Stretching stiffness for any type of bead | 0.17 kcal/mol/$\text{A}^2$ [5] |
| $k^{bend}$ | Bending stiffness of DNA bead | 8.8kcal/mol [5] |
| $\Delta t$ | Time-step for BD | $359.5fs$ |

Table 1: Parameters used in our simulations
